# Inverse Protocol Prediction from Spheroid Microscopy Imaging via Morphology-Aware Structured Learning

**DOI:** 10.64898/2026.03.04.709682

**Authors:** Prateek Mittal, Ayush Srivastava, Joohi Chauhan

## Abstract

We introduce *Inverse Protocol Prediction (IPP)*, which is a task of inferring experimental culture conditions directly from a single bright-field spheroid image. We formulate IPP as a structured multi-label prediction problem and propose a protocol-aware learning framework that integrates morphology extraction, multimodal representation learning, and dependency-aware inference. Morphometric descriptors derived from automated spheroid segmentation are fused with deep visual embeddings via a morphometry vision fusion module. To capture biological and procedural dependencies, we develop a Hierarchical Multi-Task Transformer that conditions predictions across protocol attributes. The framework is trained with domain-adversarial supervision and morphology-preserving augmentation to improve robustness to acquisition variability. Hybrid convolution attention encoders achieve the best performance, reaching 95.7% multi-attribute accuracy. We further evaluate cross dataset transfer and temporal morphology forecasting. Results demonstrate that structured, dependency-aware modeling enables reliable reconstruction of experimental protocols from imaging alone, supporting reproducibility auditing and protocol validation in 3D cell culture systems.

## Introduction and Background

Three-dimensional (3D) cell culture systems such as spheroids and organoids are central tools in cancer biology, drug discovery, and tissue engineering (1, 2). Unlike two-dimensional cultures, spheroids recreate nutrient and oxygen gradients, cell–cell interactions, and necrotic cores, while remaining amenable to bright-field and phase-contrast microscopy for high-throughput, non-destructive readouts (3). Despite this ubiquity, microscopy is still used primarily to measure experimental outcomes (e.g., size, viability, or morphology), rather than to interrogate whether the observed morphology is consistent with the experimental protocol under which it was generated.

This gap limits reproducibility and quality control: discrepancies between reported metadata (cell line, medium, seeding density, formation method, imaging setup) and the resulting morphology often remain undetected, particularly in large-scale or multi-site studies. Inverse Protocol Prediction (IPP) addresses this limitation by treating microscopy images as diagnostic signals of their underlying experimental context, enabling automated validation of protocol consistency directly from imaging data. We pose the following challenge: given a single spheroid image, can we infer the experimental protocol attributes most plausibly associated with it, such as cell line, culture medium, seeding density, timepoint, formation method, and microscope? We term this the Inverse Protocol Prediction (IPP) problem. Success would enable imagebased reproducibility checks, detection of protocol inconsistencies or labeling errors, and systematic analysis of which experimental factors leave recoverable morphological signatures in spheroid microscopy.

Recent advances in biomedical image segmentation leverage deep convolutional and transformer-based architectures to handle the visual complexity of 3D cell culture images. U-Net variants and DeepLab models achieve strong performance under noise and imaging artifacts, while transformer-based designs such as Swin-UNet capture global contextual cues and improve recognition of complex spheroid morphology (4–6). Semi-supervised learning and multi-label deep supervision address limited annotations and class imbalance, improving robustness and generalization (7, 8). Synthetic data generation guided by biophysical principles has further enhanced data diversity and realism (9). In parallel, recent work on IPP from biomedical images has adopted multilabel and transformer-based frameworks to infer experimental metadata, exploiting dependencies between protocol components and heterogeneous visual cues (10, 11). Spatiotemporal modeling using ConvLSTM and attention-based recurrent architectures has enabled time-series prediction of longitudinal morphological dynamics in microscopy data (12, 13). Incorporating metadata and multimodal inputs further improves generalization across experimental conditions.

Inverse Protocol Prediction (IPP) is difficult for several reasons. First, morphological ambiguity arises when distinct experimental conditions yield visually similar imaging outputs, and large variability in spheroid morphology remains understudied and hard to disentangle without standardized datasets linking images to detailed metadata (14, 15). Second, imaging variability across microscopes, modalities, and acquisition settings introduces substantial artifacts and inconsistencies that can obscure biologically relevant features and impede reproducibility (16, 17). Third, few microscopy datasets couple high-resolution 3D images with rich experimental metadata, limiting model development and evaluation; comprehensive reporting of imaging methods and metadata is notoriously underreported in biomedical research, fur-ther compounding this scarcity (18). To address this, we introduce a unified framework for inverse protocol prediction using SLiMIA (19), a dataset of *∼*8,000 bright-field spheroid images spanning nine microscopes, 47 cell lines, multiple media, seeding densities, timepoints, and formation methods. We frame the problem as a structured multi-label prediction, disentangle morphology from imaging artifacts via domain-adversarial training and augmentation, and benchmark convolutional, transformer, and hybrid architectures. While many components are established, our novelty lies in their adaptation to the structured multi-label nature of experimental protocols, where causal label dependencies and morphometric priors are biologically grounded rather than arbitrary.

Our main contributions are summarized as:

1. Morphometry fusion integrating classical geometric descriptors with deep visual embeddings for protocol-aware feature learning.
2. Hierarchical multi-task transformer modeling for structured Inverse Protocol Prediction (IPP).
3. Protocol-level interpretability via Grad-CAM, analyzing biological versus acquisition-driven prediction signals.
4. Spatiotemporal extension of IPP through time-series prediction of longitudinal spheroid evolution using ConvLSTM, PredRNN++, and PhyDNet.
5. Systematic evaluation of protocol attribute recoverability, revealing which experimental factors leave stable morphological signatures.

This work demonstrates that spheroid morphology encodes rich, recoverable signatures of culture conditions, establishing microscopy-driven inverse protocol prediction as a new paradigm for reproducibility, optimization, and design in cell culture systems.

## Methodology and Experimental Results

### Dataset Description

SLiMIA (Spheroid Light Microscopy Image Atlas) (19) is an open-access morphometric image dataset designed to support machine learning and computational modeling of three-dimensional (3D) cell culture systems. SLiMIA comprises approximately 8,000 light microscopy images of spheroids spanning a diverse range of experimental conditions, including nine microscope types, 47 distinct cell lines, eight culture media, four spheroid formation protocols, and multiple cell seeding densities, accompanied by rich metadata to facilitate reproducible analysis and benchmarking. Figure 1 highlights some sample dataset images along wih their manual segmentatons. Figure 2 demonstrates the entire workflow followed for analysis of the dataset. The same is described in detail in this section.

**Fig. 1.**
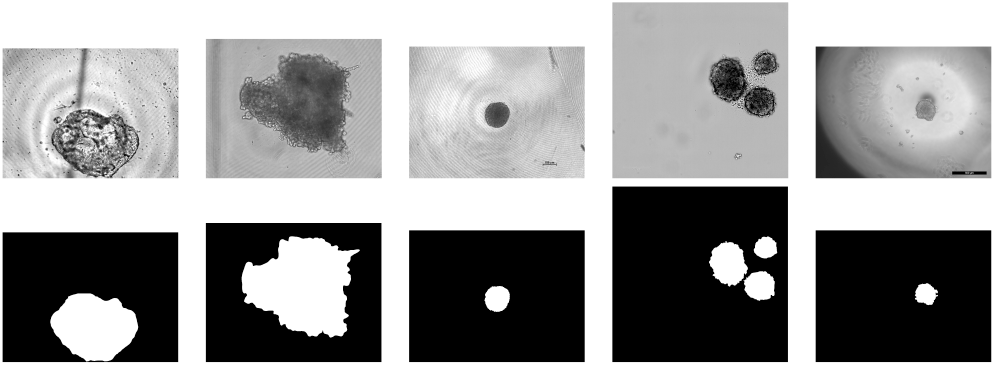
Sample images from the SLiMIA dataset with their manual segmentations (bottom row).

**Fig. 2.**
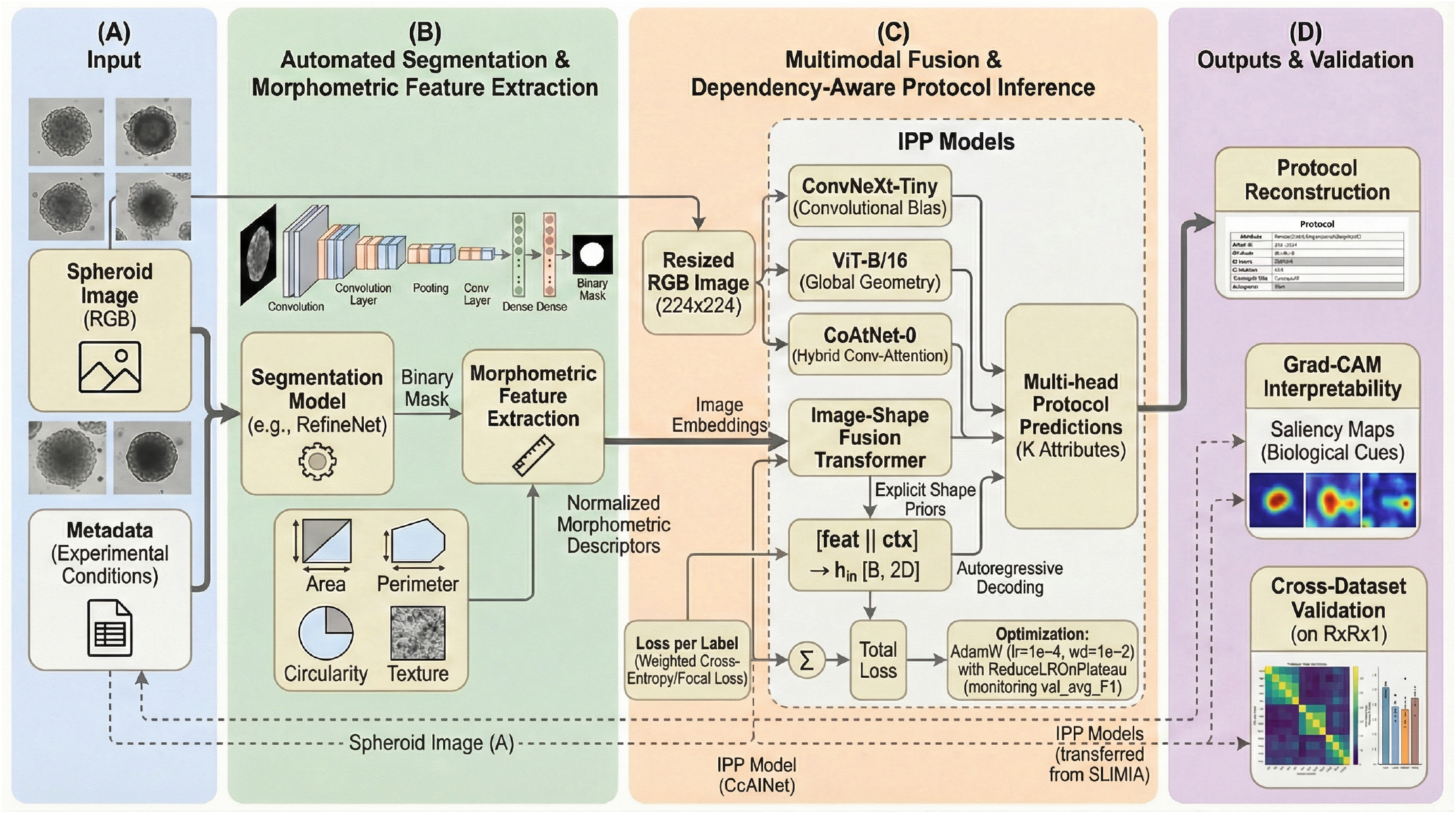
Inverse protocol prediction (IPP) workflow. (A) Input spheroid image and metadata. (B) Automated segmentation and morphometric feature extraction. (C) Multimodal fusion and dependency-aware protocol inference using HMTT (D) Outputs including protocol reconstruction, Grad-CAM interpretability and cross data validation

**Fig. 3.**
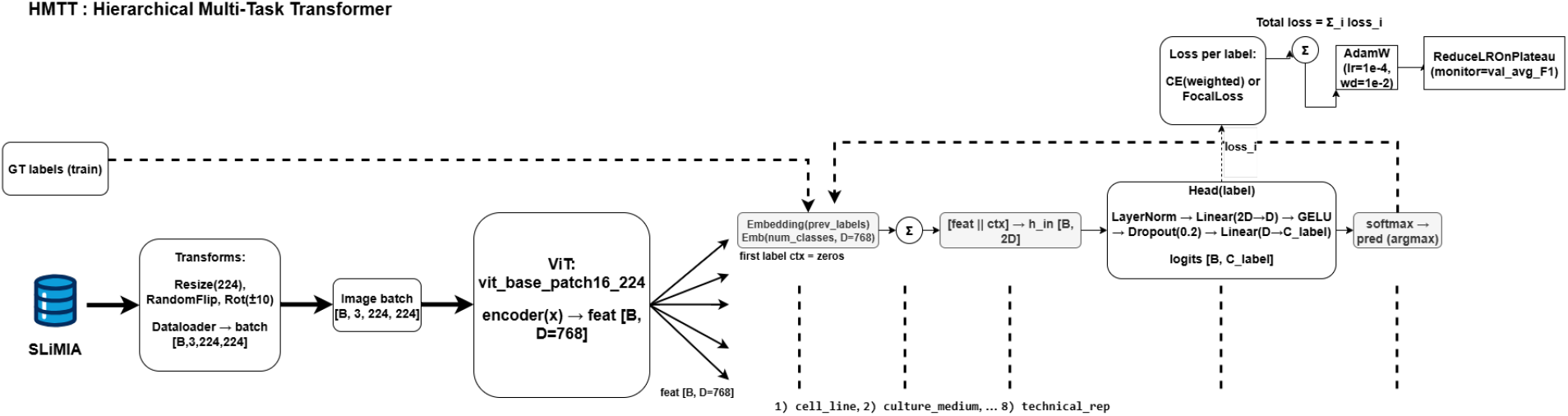
Overview of the Hierarchial Multi Task Transformer (HMTT) architecture

### Train–Validation Partition Strategy

To prevent image-level leakage across acquisition sessions, data were partitioned at the level of technical replicate identifiers (T1–T24). All images belonging to a given technical replicate were grouped prior to splitting to ensure that no acquisition session appeared in more than one subset.

Technical replicates were manually distributed across training, validation, and test partitions to balance sample size while maintaining strict replicate exclusivity. Replicates T1–T4 were assigned to the training set, T5 and T8 were assigned to the validation set, and T6, T7, T9–T24 were reserved exclusively for testing. This allocation ensures that each split contains multiple acquisition sessions, avoids single-replicate validation bias, and distributes larger replicate groups across partitions to stabilize optimization and evaluation. No random image-level splitting across technical replicate identifiers was performed. Consequently, images originating from a given acquisition session do not appear in multiple subsets, preventing acquisition-specific information leakage.

Final model performance was evaluated exclusively on the held-out test partition, which contains technical replicates not seen during training or validation. This design provides a conservative estimate of biological attribute inference under cross-replicate acquisition variability.

### Attribute-Level Class Coverage Analysis

Following replicate-level partitioning, we conducted an explicit audit of class coverage across the training, validation, and test splits for each metadata attribute to quantify potential distribution shift. For each attribute, we computed the total number of unique classes in the full dataset and verified the number of classes represented within each split.

Across seven of the eight attributes (microscope, cell line, culture medium, formation method, seeding density, biological rep, and magnification), all classes observed in the validation and test sets were also present in the training set. This confirms the absence of unseen attribute categories at evaluation time for these metadata variables and indicates that performance estimates reflect generalization across acquisition sessions rather than exposure to novel class identities.

For the timepoint attribute (109 total classes), 107 classes were represented in the training partition. Two classes (000h and 032h) were present in the test set but absent from training. No unseen validation classes were observed relative to the training set. This limited exposure to unseen timepoint categories represents a minor open-set condition affecting a small subset of samples.

Overall, the dataset exhibits near-complete attribute coverage across splits, with strict technical replicate exclusivity preserved and minimal unseen-category presence confined to a small subset of timepoint labels. This analysis confirms that the evaluation protocol primarily measures cross-replicate generalization under acquisition variability rather than robustness to large-scale unseen class distributions.

### Segmentation Models

All segmentation models were trained on RGB inputs with a single binary mask output. CNN backbones were initialized with ImageNet pretraining. Optimization was performed using Adam or AdamW (20, 21) with ReduceLROnPlateau scheduling (patience = 5, factor = 0.5) (22). Loss functions were selected based on architectural characteristics, balancing region overlap and class imbalance (e.g., Focal Tversky (23) or BCE+Dice (24)). Standard data augmentations including flips and intensity/geometric perturbations were applied during training. Backbone encoders and training configurations are summarized in Table 1.

**Table 1.** Segmentation model configurations and training settings. All CNN backbones were initialized with ImageNet pretraining.

| Model | Backbone | Loss | Optimizer |
| --- | --- | --- | --- |
| U-Net++ (25) | ResNet-50 | Focal Tversky (23) ($\alpha=0.7$, $\beta=0.3$, $\gamma=0.75$) | Adam (20) + ReduceLROnPlateau (22) |
| DeepLabV3+ (26) | ResNet-50 | Focal Tversky (23) | Adam (20) + ReduceLROnPlateau (22) |
| DeepLabV3 (26) | ResNet-34 | BCE + Dice (24) | AdamW (21) (3e-4, wd=1e-4) |
| Attention U-Net (27) | ResNet-based | BCE + Dice (24) | AdamW (21) |
| Swin-UNet (28) | Swin-Tiny | BCEWithLogits (29) | Adam (20) (1e-4) |
| TransUNet (30) | ViT Encoder + CNN Decoder | BCEWithLogits (29) | Adam (20) (1e-4) |
| RefineNet (31) | ResNet-34 | Focal Tversky (23) | Adam (20) + ReduceLROnPlateau (22) |
| SegNet (32) | VGG-style Encoder-Decoder | Focal Tversky (23) | Adam (20) + ReduceLROnPlateau (22) |

**Table 2.**
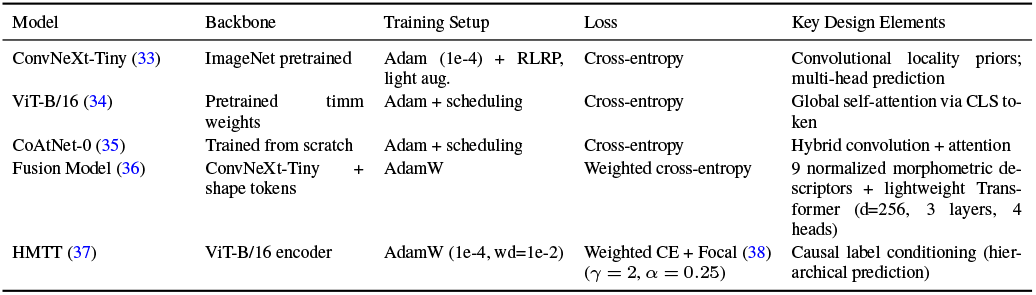
Implementation details of IPP models.

Morphometric descriptors used in downstream IPP models are computed exclusively from automatically predicted segmentation masks. A dedicated segmentation network (RefineNet; best-performing model in Table 1) is first trained using annotated masks and subsequently frozen. Predicted masks are then used to extract quantitative morphometric features for all IPP experiments. Ground-truth masks are never used during IPP training or evaluation, ensuring a fully automatic pipeline and preventing annotation leakage.

### Implementation Details of Inverse Protocol Prediction (IPP) Models

We formulate IPP as a structured multi-label prediction task, predicting experimental attributes from a single spheroid image. All models use RGB inputs resized to 224×224 and a multi-head design with one classifier per protocol label.

ConvNeXt-Tiny (33) serves as a strong convolutional baseline, preserving locality priors beneficial for fine-grained morphological cues such as magnification and seeding density. ViT-B/16 (34) emphasizes global self-attention, capturing long-range dependencies relevant for formation method and temporal progression. CoAtNet-0 (35) combines convolution and attention, balancing texture sensitivity and structural modeling. To incorporate explicit morphometric priors, we introduce a fusion model that augments ConvNeXt embeddings with normalized shape descriptors, enabling improved robustness and interpretability through joint feature learning (36). Finally, the Hierarchical Multi-Task Transformer (HMTT) enforces a predefined causal ordering among protocol components, sequentially conditioning each attribute prediction on previously inferred labels to promote biologically coherent reconstructions (37). The hierarchical order is specified a priori based on experimental protocol structure, progressing from stable upstream determinants (e.g., cell line and culture medium) to condition-dependent variables (e.g., seeding density and timepoint), and is implemented using teacher-forced conditioning during training and autoregressive inference at evaluation time. This design reflects the causal progression of spheroid formation while acknowledging that early prediction errors may propagate to downstream attributes under autoregressive decoding. In addition to architectural conditioning, we employ conservative morphology-preserving data augmentation during IPP training to enhance invariance without distorting biologically meaningful structure. Specifically, input images are resized to a fixed spatial resolution and augmented using small-angle rotations ( *±*10^*°*^) and horizontal flipping, followed by normalization. These transformations enforce orientation robustness while preserving spheroid topology, boundary integrity, and internal texture patterns. No elastic deformations, random cropping, scale jitter, color perturbations, or intensity-based augmentations were applied, as such operations could alter morphometric cues linked to seeding density or developmental stage. While spatial resizing standardizes input dimensions and may attenuate absolute size cues relevant to density estimation, timepoint-dependent features such as compactness and necrotic-core formation remain structurally intact. This restrained augmentation strategy ensures improved generalization across acquisition conditions without introducing artificial morphological artifacts or confounding biologically grounded attribute predictions. Together, these models span a spectrum from purely convolutional to purely transformer-based, hybrid, feature-augmented, and dependency-aware architectures. This systematic design allows us to disentangle the contributions of inductive bias, explicit priors, and label-structure modeling to the inverse inference task.

### Domain-Adversarial Training

To reduce acquisition-specific bias, we incorporated domain-adversarial training into the best-performing backbone (CoAtNet-0). The domain variable was defined as the *technical replicate identifier* (T1– T24), representing acquisition sessions.

The architecture consists of three components:

**Feature extractor:** A CoAtNet-0 backbone producing a global feature representation.

**Biological prediction heads:** Independent linear classifiers for each biological attribute (cell line, culture medium, formation method, seeding density, timepoint, biological replicate, magnification).

**Domain discriminator:** A two-layer multilayer perceptron attached through a Gradient Reversal Layer (GRL):

- Linear (*d →* 256)
- ReLU activation
- Dropout (*p* = 0.3)
- Linear (256 *→ N*_*domain*_)

The GRL acts as identity during forward propagation and multiplies gradients by *−λ* during backpropagation, encouraging domain-invariant feature learning.

The adversarial coefficient *λ* follows the standard DANN schedule:

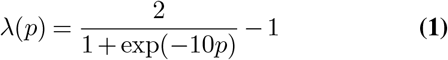

where *p∈* [0, 1] denotes normalized training progress. The total training objective is:

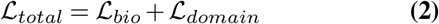

where both terms are standard cross-entropy losses and are equally weighted.

### Implementation Details of Time-Series Prediction Models

Temporal sequences were constructed by grouping SLiMIA images under consistent experimental conditions and sorting by time. Two consecutive frames were used as input to predict the subsequent frame, with variable temporal gaps (1–3 intervals) to increase diversity and capture short- and mid-range dynamics. Implementation details of all models are summarized in Table 3.

**Table 3.** Implementation details of time-series prediction models. All models use two input frames with variable temporal gaps (1–3 intervals) for next-frame prediction.

| Model | Architecture | Training Setup | Loss | Key Design Elements |
| --- | --- | --- | --- | --- |
| ConvLSTM (39) | Single ConvLSTM (32 ch.) + Conv decoder | Adam, variable gap (1–3) | L1 | Convolutional recurrent modeling of spatial–temporal correlations |
| PredRNN++ (40) | 4-layer ST-LSTM (32,64,64,64) + GHU | Adam, variable gap (1–3) | L1 | Dual-memory spatiotemporal recurrence with gradient high-way unit |
| MetadataFusion (41, 42) | CNN encoder–decoder + metadata embeddings | AdamW | MSE + L1 | FiLM conditioning (43) using categorical embeddings and continuous MLP features |
| PhyDNet (44, 45) | PhyCell + residual ConvLSTM | Adam + L1 regularization on physics operator | L1 | Physics-guided recurrent cell + residual dynamics; FiLM conditioning |

In total, 5,770 unique experimental groups were identified from the metadata. After enforcing the requirement of at least two input frames and one future target frame, 616 groups contained sufficient longitudinal observations for temporal modeling. Using a sequence length of *L* = 2 and enumerating all admissible sliding windows with temporal gaps Δ*t ∈ {* 1, 2, 3 *}* relative to the final input frame, a total of 3,015 input–target sequences were generated for training and evaluation. Grouping was performed across fixed metadata attributes including microscope, cell line, culture medium, formation method, seeding density, magnification, biological replicate, and technical replicate to ensure condition consistency within each temporal trajectory.

### Results and Discussion

We benchmarked eight segmentation architectures spanning classical CNN encoder–decoders, refinement-focused designs, and transformer-based models to evaluate robustness on spheroid microscopy images. The selected models address key challenges including faint boundaries, class imbalance, and intensity variability. CNN-based baselines (U-Net++, Attention U-Net, SegNet, RefineNet) represent varying design priorities, from lightweight efficiency to boundary refinement. DeepLabV3/V3+ provide strong atrous convolution baselines with powerful backbones. Transformer-based Swin-UNet and TransUNet capture long-range contextual dependencies via self-attention. Together, these architectures offer a balanced comparison across convolutional and attention-driven paradigms. Quantitative results are summarized in Table 4, with qualitative examples shown in Figure 4.

**Table 4.** Segmentation performance on the SLiMIA dataset. Metrics are reported as mean ± standard deviation computed across test images. 95% confidence intervals were derived as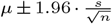.

| Model Name | Dice | IoU | Accuracy | Precision | Recall | F1 | Inf. Time (s) |
| --- | --- | --- | --- | --- | --- | --- | --- |
| U-Net++ (25) | 0.9582 ± 0.00019 | 0.9316 ± 0.00023 | 0.9929 ± 0.00002 | 0.9737 ± 0.00016 | 0.9447 ± 0.00018 | 0.9582 ± 0.00019 | 262 |
| SegNet (32) | 0.9521 ± 0.00020 | 0.9178 ± 0.00025 | 0.9913 ± 0.00001 | 0.9666 ± 0.00017 | 0.9402 ± 0.00019 | 0.9521 ± 0.00020 | 173 |
| Swin-UNet (28) | 0.9467 ± 0.00021 | 0.9103 ± 0.00026 | 0.9907 ± 0.00002 | 0.9428 ± 0.00016 | 0.9524 ± 0.00018 | 0.9467 ± 0.00021 | 208 |
| TransUNet (30) | 0.9579 ± 0.00019 | 0.9260 ± 0.00023 | 0.9929 ± 0.00002 | 0.9618 ± 0.00016 | 0.9554 ± 0.00018 | 0.9579 ± 0.00019 | **164** |
| Attention U-Net (27) | <u>0.9604 ± 0.00018</u> | <u>0.9361 ± 0.00022</u> | <u>0.9934 ± 0.00002</u> | 0.9615 ± 0.00017 | **0.9609 ± 0.00018** | <u>0.9604 ± 0.00018</u> | <u>166</u> |
| DeepLabV3 (46) | 0.9571 ± 0.00019 | 0.9302 ± 0.00023 | 0.9928 ± 0.00003 | 0.9596 ± 0.00016 | 0.9568 ± 0.00018 | 0.9571 ± 0.00019 | 184 |
| DeepLabV3+ (46) | 0.9551 ± 0.00019 | 0.9271 ± 0.00023 | 0.9925 ± 0.00002 | 0.9710 ± 0.00016 | 0.9426 ± 0.00020 | 0.9551 ± 0.00019 | 239 |
| RefineNet (31) | **0.9665 ± 0.00016** | **0.9437 ± 0.00021** | **0.9938 ± 0.00001** | **0.9765 ± 0.00015** | <u>0.9576 ± 0.00018</u> | **0.9665 ± 0.00016** | 286 |

**Fig. 4.**
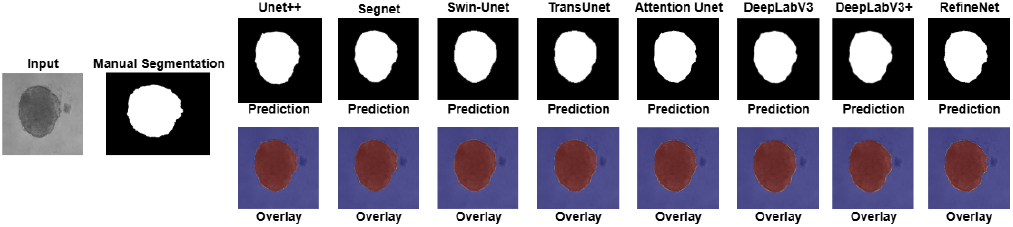
Comparison of segmentation results across different segmentation models. The first column shows the input image and corresponding manual segmentation (ground truth). Subsequent columns display predictions and overlay results for U-Net++, SegNet, Swin-Unet, TransUnet, Attention U-Net, DeepLabV3, DeepLabV3+, and RefineNet.

We calculated the average Dice and Intersection over Union (IoU) scores across all images in the dataset, while explicitly excluding images for which the Dice score equals 1 and the IoU score equals 0. These cases represent edge scenarios where both predicted and ground truth masks are empty, leading to metric inconsistencies. Moreover, among the dataset, 17 such images have corresponding masks unavailable. By excluding these from the metric averages, we ensure a more accurate and meaningful assessment of segmentation performance on images containing valid target structures.

Overall, all architectures performed strongly with Dice scores above 0.94, confirming the suitability of modern segmentation models for spheroid analysis. RefineNet achieved the best results, likely due to its refinement blocks that focus on structural boundaries and multi-scale context, beneficial for faint or irregular spheroid edges, though it had the longest inference time. Attention U-Net and U-Net++ performed competitively, balancing accuracy and moderate inference time, helped by skip connections and attention that capture fine details and reduce background noise. Transformer-based models like Swin-UNet and TransUNet had slightly lower scores, probably due to limited data and their data-intensive nature, but were more efficient in inference than deeper CNNs. SegNet performed respectably with one of the fastest inference times, showing that classical encoder–decoders remain viable baselines. These results highlight the advantage of boundary-focused designs for spheroid segmentation, trade-offs between lightweight CNNs and refinement-heavy models in efficiency, and the need for larger datasets or pretraining to fully exploit transformer-based methods in biomedical imaging.

### A. Evaluation of Inverse Protocol Prediction (IPP) Models

IPP reconstructs experimental conditions from spheroid morphology, requiring models that capture fine-grained local texture, global structural context, and dependencies among protocol labels. We evaluate five complementary architectures (Table 5) to probe these requirements. ConvNeXt-Tiny serves as a modern convolutional baseline with strong locality biases, while ViT-B/16 models long-range spatial interactions critical for global spheroid geometry. CoAtNet bridges these paradigms through hybrid convolution–attention blocks. Beyond generic backbones, we introduce two spheroid-specific designs: an Image–Shape Fusion Transformer that injects explicit morphometric priors for improved robustness and interpretability, and a Hierarchical Multi-Task Transformer (HMTT) that enforces biologically motivated label dependencies to produce causally consistent protocol predictions. Together, these models isolate the roles of inductive bias, explicit priors, and hierarchical structure in inverse protocol inference from microscopy images.

**Table 5.**
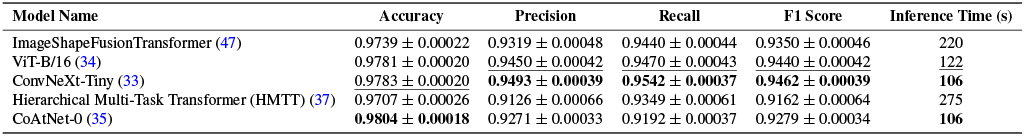
IPP model performance reported as mean *±* standard deviation across three independent random seeds (42, 123, 999). Model-specific 95% confidence intervals are computed using the normal approximation (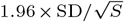, with *√S* = 3 seeds).

#### Definition of Multi-Attribute Accuracy

IPP is formulated as a multi-label multi-class prediction problem with *K* protocol attributes per image. We report two complementary definitions of multi-attribute accuracy:

#### (1) Exact-Match Accuracy (Subset Accuracy)

An image is considered correct only if *all K* predicted attributes match the ground truth:

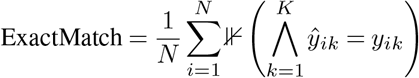

where *N* is the number of test images and *K* the number of attributes.

#### (2) Mean Per-Attribute Accuracy

We also compute the average accuracy across individual attribute heads:

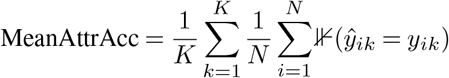

Unless otherwise stated, the accuracy reported in Table 5 corresponds to Mean Per-Attribute Accuracy. Exact-match results are reported separately below.

Overall, the architectures exhibit complementary strengths across biological protocol variables. CoAtNet achieved the highest macro-averaged accuracy (98.0%), indicating strong overall classification reliability. However, ConvNeXt-Tiny achieved the highest macro F1 score (0.946), driven by superior precision and recall balance across labels. ViT-B/16 performed comparably, showing consistent recall and competitive F1 performance. The Image–Shape Fusion Transformer remained competitive (macro F1 = 0.935), supporting the value of incorporating explicit morphometric priors, though at increased computational cost. While HMTT achieved lower macro accuracy (97.1%) and F1 (0.916), its hierarchical formulation yielded stable performance across interdependent labels. Per-label analysis (Table 6, Appendix A.1) shows that attributes with strong morphological signatures (cell line, medium, formation method) are predicted reliably, whereas weaker or less visually encoded factors (seeding density, timepoint, replicate) remain challenging. Near-perfect microscope and magnification scores likely reflect dataset-specific artifacts rather than intrinsic biological information.

**Table 6.** Per-label best performance across models. Labels are grouped into biological protocol variables and acquisition/replicate variables. The Biological Macro F1 (0.915) is computed over cell line, culture medium, seeding density, timepoint, and formation method only, isolating biologically meaningful inference from acquisition-specific cues. Technical replicate identifiers are excluded from macro-level aggregation, as data partitioning was performed at the replicate level to prevent information leakage across acquisition sessions.

| Label | Best F1 | Best Model | Average F1 (all models) |
| --- | --- | --- | --- |
| *Biological Protocol Variables* |  |  |  |
| Cell Line | 0.9944 | ImageShapeFusionTransformer | 0.9888 |
| Culture Medium | 0.9642 | ViT | 0.9604 |
| Seeding Density | 0.9302 | CoAtNet | 0.8871 |
| Timepoint | 0.8331 | ConvNeXt | 0.7494 |
| Formation Method | 0.9949 | ConvNeXt | 0.9940 |
| **Biological Macro F1** | – | – | **0.915** |
| *Acquisition and Replicate Variables* |  |  |  |
| Biological Rep | 0.9583 | ConvNeXt | 0.9444 |
| Magnification | 0.9997 | CoAtNet | 0.9985 |
| Microscope | 1.0000 | ImageShapeFusionTransformer | 0.9993 |

#### Biological-Only Inverse Protocol Prediction

To assess protocol inference independently of acquisition-specific cues, we recomputed IPP performance after excluding microscope, magnification, and replicate labels. The resulting biological-only benchmark includes five attributes: cell line, culture medium, seeding density, timepoint, and formation method (Table 6).

Across models, the macro-averaged Biological IPP F1 score is 0.915. High recoverability is observed for cell line (F1 *≈*0.99) and formation method (F1 *≈*0.99), indicating that these protocol variables produce consistent and discriminative morphological patterns. Culture medium and seeding density remain reliably inferable, though increased visual similarity between classes reduces separability. Timepoint prediction is comparatively more challenging (F1 *≈*0.75), reflecting subtle morphological differences between adjacent developmental stages.

Importantly, strong performance persists even after excluding acquisition-driven variables, supporting the presence of biologically meaningful structural variation encoded in spheroid morphology.

#### A.1. Effect of Domain-Adversarial Training

We evaluated biological prediction and domain predictability on the held-out test set.

#### Biological performance (CoAtNet + DANN)

Macro (across heads): Accuracy = 0.8973, Precision = 0.8267, Recall = 0.7979, F1 = 0.8010.

Micro (flattened across attributes): Accuracy / F1 = 0.8973.

### B. Interpretability Analysis with Grad-CAM

To examine the internal reasoning of our IPP framework, we applied Grad-CAM analysis (48, 49) to the best-performing architecture, CoAtNet. Its hybrid convolution–attention design is well-suited to SLiMIA, where both local morphology and global context are critical. The goal was to verify reliance on biologically meaningful cues and identify potential dataset-specific biases.

We analyzed ten representative spheroids spanning diverse cell lines (A549, HCT116, PANC1, SKOV3, U251MG, SW837, MCF10A, CT5.3hTERT), media, formation protocols (ULA, Hanging Drop, Agarose, Microchip), seeding densities, timepoints, replicates, and magnifications, ensuring broad experimental coverage.

Grad-CAM results (Figure 5) show consistent attention to biologically relevant features. For cell line prediction, attention concentrates on global morphology and boundary structure. Formation method and seeding density focus on compactness and internal organization, while later timepoints emphasize dense cores consistent with spheroid maturation. In contrast, replicate-related predictions yield diffuse or background-oriented attention, suggesting sensitivity to experimental artifacts. The model also adapts attention across magnifications, highlighting global structure at low resolution and finer boundary detail at higher resolution.

**Fig. 5.**
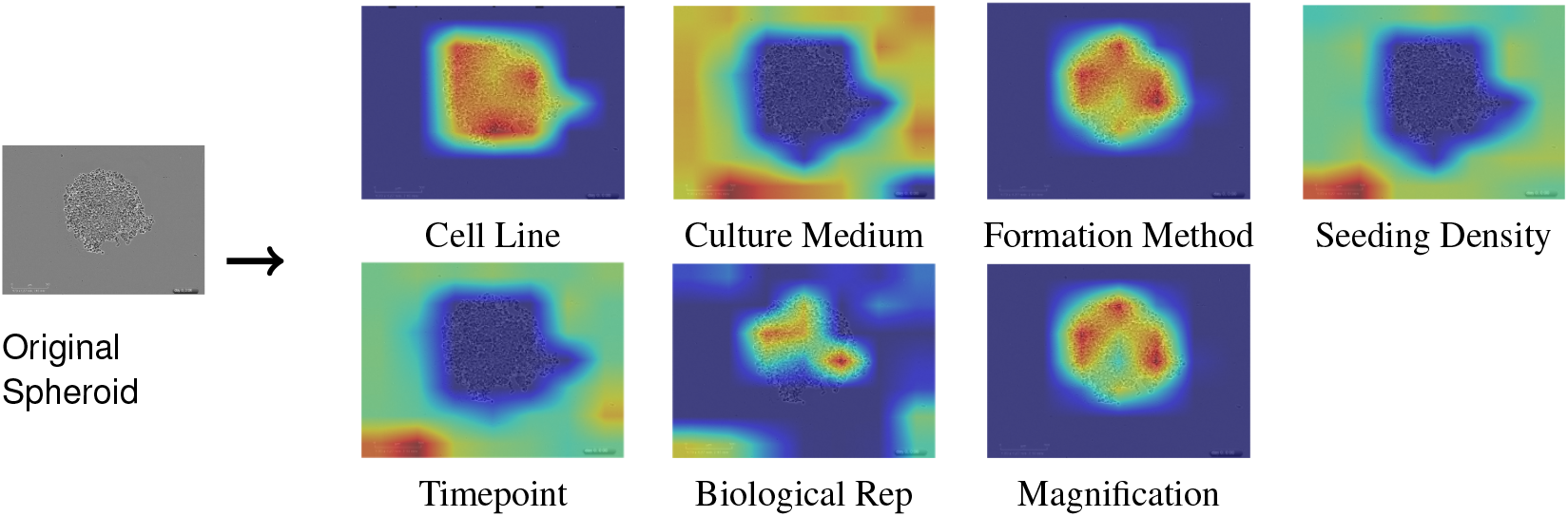
Grad-CAM visualizations for inverse protocol prediction. Left: original spheroid image. Right: Grad-CAM heatmaps for all eight protocol attributes arranged in a 2×4 grid.

Overall, Grad-CAM confirms that CoAtNet primarily relies on interpretable morphological cues for key biological variables, while revealing vulnerability to technical confounders in replicate-level predictions. These findings support the architectural suitability of CoAtNet for IPP and underscore the importance of interpretability in distinguishing biological reasoning from dataset-specific artifacts.

### C. Evaluation of Time Series Prediction Models

To model spheroid growth dynamics, we evaluated ConvLSTM and PredRNN++ as recurrent baselines. ConvLSTM incorporates convolutional gates to capture spatial–temporal correlations(39), while PredRNN++ introduces dual-memory mechanisms for improved long-term dependency modeling(40). To assess contextual conditioning, we included a Metadata Fusion model that integrates image features with structured protocol metadata(41, 42), and PhyDNet, which separates physical dynamics from residual components to better reflect biological processes(44). Results are summarized in Table 7.

**Table 7.** Spatiotemporal prediction performance with model-specific 95% confidence intervals. Confidence intervals reflect sample-level variability estimated using the normal approximation 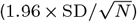 with *N ≈* 8000.

**Table 8.**
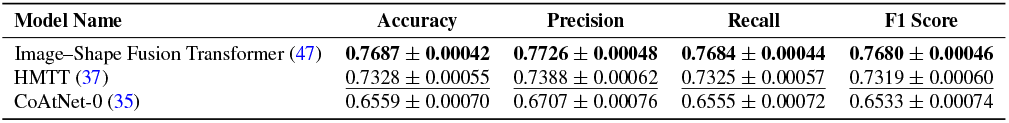
Cross-dataset IPP results on RxRx1 with 95% confidence intervals.

**Table 9.** Cross-dataset temporal validation on the CTC dataset.

| Model | MSE | SSIM | PSNR (dB) |
| --- | --- | --- | --- |
| PredRNN++ | $0.002753 \pm 0.000033$ | $0.5903 \pm 0.0021$ | $25.60 \pm 0.033$ |
| ConvLSTM | $0.003763 \pm 0.000036$ | $0.5154 \pm 0.0022$ | $24.24 \pm 0.034$ |

The results (Figure 6) indicate modest performance overall (SSIM *<* 0.40, PSNR *≈*18 dB). MetadataFusion performs best, demonstrating the benefit of protocol-aware conditioning. PhyDNet improves structure modeling through physics-inspired constraints. However, temporal prediction remains challenging due to non-linear growth processes (pro-liferation, compaction, necrosis) and short, irregular SLiMIA sequences, limiting long-term dependency learning. These findings suggest that richer longitudinal data or hybrid mechanistic models are needed for improved forecasting.

**Fig. 6.**
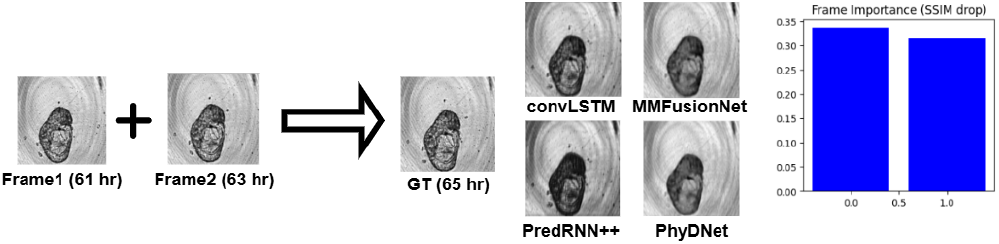
Time-series prediction results. Two consecutive input frames (61 hr, 63 hr) are used to predict the next frame (65 hr) by four different models, compared against the ground truth.A frame importance plot shows how omitting each input frame affects SSIM, highlighting their relative contribution.

**Fig. 7.**
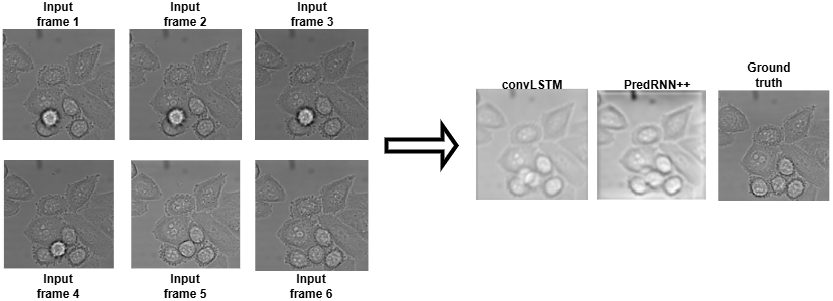
Temporal prediction on the CTC dataset. One observed frame plus five subsequent frames are used to predict the future frame. ConvLSTM and PredRNN++ outputs are shown alongside ground truth.

### D. Statistical Analysis

To evaluate whether CoAtNet exhibits consistent per-attribute performance differences relative to competing architectures, we conducted paired two-sided statistical tests on F1 scores across eight biologically meaningful labels (Microscope, Cell Line, Culture Medium, Formation Method, Seeding Density, Timepoint, Biological Rep, Magnification). Each comparison therefore used *n* = 8 paired observations (*df* = 7).

For each model comparison, the null hypothesis was:

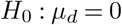

where *µ*_*d*_ denotes the mean paired difference in F1 score between CoAtNet and the comparison model. The alternative hypothesis was:

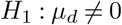

Statistical significance was evaluated at *α* = 0.05. In addition to paired *t*-tests, Wilcoxon signed-rank tests were performed to mitigate normality assumptions. Effect sizes were quantified using Cohen’s *d*_*z*_, and 95% confidence intervals were computed for the mean differences. Holm–Bonferroni correction was applied across the four pairwise comparisons.

#### CoAtNet vs ConvNeXt-Tiny

No statistically significant difference was observed (*t*(7) = *−*0.84, *p* = 0.43, *d*_*z*_ = *−*0.30, 95% CI [ *−*0.095, 0.045]). The Wilcoxon test yielded *p* = 0.46.

#### CoAtNet vs HMTT

No significant difference was detected (*t*(7) = 0.19, *p* = 0.86, *d*_*z*_ = 0.07, 95% CI [ *−*0.028, 0.033]). Wilcoxon testing produced *p* = 0.88.

#### CoAtNet vs ImageShapeFusion

The comparison did not reach statistical significance (*t*(7) = *−*1.45, *p* = 0.19, *d*_*z*_ = *−*0.51, 95% CI [ *−*0.061, 0.015]). The Wilcoxon test yielded *p* = 0.22.

#### CoAtNet vs ViT

No statistically significant difference was observed (*t*(7) = *−*1.01, *p* = 0.34, *d*_*z*_ = *−*0.36, 95% CI [*−*0.100, 0.040]). Wilcoxon testing produced *p* = 0.36.

#### Interpretation

Across eight attribute-level evaluations, no comparison demonstrated statistically significant differences after Holm–Bonferroni correction. All confidence intervals included zero, indicating absence of uniform per-attribute superiority. Variability was primarily driven by the Timepoint attribute, suggesting that performance differences are concentrated in specific protocol variables rather than representing a consistent global shift across biological factors.

### Cross-Dataset Generalization Analysis

#### Cross-Dataset Validation on RxRx1 for Protocol Prediction (IPP)

To rigorously evaluate cross domain generalization of the Inverse Protocol Prediction (IPP) framework, we conducted validation on the RxRx1 dataset (50). RxRx1 is a large scale fluorescence microscopy benchmark comprising chemically perturbed monolayer cells collected across multiple experimental batches and imaging sites, exhibiting strong batch effects and substantial morphological variability. In contrast to SLiMIA’s 3D spheroid structures, RxRx1 contains 2D monolayer cell populations. For RxRx1, we adapt the IPP formulation to predict experimental metadata attributes directly from fluorescence microscopy images: *experiment* (51 classes), *plate* (4 classes), *site* (2 classes), *cell type* (4 classes), and *sirna* (1139 classes), rather than the standard compound identification task. This extends the inverse protocol prediction framework to a new experimental context by inferring acquisition and experiment level metadata directly from morphology. The high cardinality of the *sirna* attribute in particular introduces a substantially more challenging multi class setting relative to SLiMIA.

The corresponding uniform chance level baselines given by 1*/K* are 0.0196 for experiment, 0.25 for plate, 0.50 for site, 0.25 for cell_type, and 0.000878 for sirna. Majority class baselines are similarly low for most attributes, namely 0.0196 for experiment, 0.2500 for plate, 0.50 for site, and 0.00392 for sirna, with the exception of *cell_type* at 0.470, reflecting moderate class imbalance. Under independent uniform guessing, the exact match subset baseline for simultaneously predicting all five attributes is 5.38 *×*10^*−*7^, while the majority class exact match baseline is 4.53*×* 10^*−*6^, underscoring that correct multi attribute reconstruction constitutes a highly non trivial objective.

This structural mismatch introduces a severe domain shift in geometry, texture statistics, and acquisition artifacts, making it a stringent test of IPP robustness. We used 125,511 Channel 1 images and constructed a strict 70:15:15 train validation test split grouped by `sirna_id` (Train 87,857, Validation 18,826, Test 18,827). Group wise splitting prevents label leakage and avoids near duplicate morphological patterns appearing across partitions, ensuring a fair evaluation of generalization. Importantly, models were transferred directly from SLiMIA without fine tuning, isolating representation robustness rather than adaptation capacity.

We evaluated three top-performing SLiMIA models: Image–Shape Fusion Transformer, Hierarchical Multi-Task Transformer (HMTT), and CoAtNet-0. Confidence intervals were computed using the normal approximation 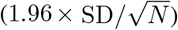 over the full RxRx1 test partition (*N* = 18, 827).

The Image–Shape Fusion Transformer achieves the strongest cross-domain performance, suggesting that integrating explicit morphometric priors improves invariance to structural shifts between 3D spheroids and 2D monolayers. HMTT remains competitive, indicating that modeling structured label dependencies confers some robustness; however, its hierarchical constraints appear more sensitive to perturbation-induced visual variability. CoAtNet-0 exhibits the largest performance degradation, consistent with reliance on spatial inductive biases tuned to spheroid geometry.

Overall, these findings indicate that feature-augmented and dependency-aware architectures generalize more effectively under severe geometric and acquisition shifts. The results support the hypothesis that incorporating explicit priors and structured prediction mechanisms enhances robustness beyond purely convolutional or hybrid attention backbones.

#### Cross-Dataset Temporal Validation on the Cell Tracking Challenge (CTC) for Time Series Prediction

We further evaluated temporal generalization using the Cell Tracking Challenge dataset (51), which provides challenging microscopy sequences with heterogeneous cell morphologies and acquisition conditions.

PredRNN++ demonstrates superior temporal generalization, achieving higher SSIM (+14.5%) and PSNR (+1.36 dB) than ConvLSTM. Its spatiotemporal memory design enables better long-range dependency retention, whereas ConvLSTM exhibits blur under strong domain shift.

### Ablation Study

To assess the contribution of each component of our fusion architecture, we conducted a series of ablation experiments. Five model variants were evaluated: (i) the full multimodal fusion model, (ii) an image-only model that removes all morphology (shape) features, (iii) a shape-only model that excludes image information, (iv) a no-fusion model where image and morphology pathways are present but not interactively fused, and (v) a no-transformer variant that retains both modalities but removes the transformer-based fusion mechanism. All models were trained under identical conditions, using the same dataset splits, augmentation strategy, and early stopping criteria to ensure a fair comparison. Expanded per-attribute ablation metrics are provided in Appendix X.

Table 10 summarizes the performance of each variant under identical IPP conditions, using the same biological-only label setting, data splits, and macro-F1 evaluation protocol. We report average accuracy, precision, recall, and macro-F1 across the biological protocol attributes. The full model achieves the strongest performance (macro-F1 = 0.8681), while the image-only model performs slightly worse (macro-F1 = 0.8662). The modest improvement (ΔF1 = 0.0019) suggests that modern convolutional backbones such as ConvNeXt already encode substantial shape, compactness, and texture information during training. Explicit morphometric descriptors therefore provide complementary structural refinement but add limited marginal value beyond deep visual representations learned directly from microscopy images.

**Table 10.** Ablation study results for multimodal metadata prediction. Values show mean accuracy, precision, recall, and macro-F1 score with estimated 95% confidence intervals computed using the normal approximation (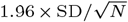, with *N ≈* 8000 validation samples).

| Model Variant | Accuracy | Precision | Recall | Macro-F1 |
| --- | --- | --- | --- | --- |
| Full Model (Image + Shape + Transformer Fusion) | $0.8526 \pm 0.00042$ | $0.8672 \pm 0.00046$ | $0.8758 \pm 0.00044$ | $0.8681 \pm 0.00045$ |
| Image-Only | $0.8516 \pm 0.00045$ | $0.8651 \pm 0.00049$ | $0.8734 \pm 0.00047$ | $0.8662 \pm 0.00048$ |
| No-Transformer | $0.8513 \pm 0.00047$ | $0.8638 \pm 0.00050$ | $0.8726 \pm 0.00049$ | $0.8650 \pm 0.00049$ |
| No-Fusion | $0.8478 \pm 0.00053$ | $0.8597 \pm 0.00056$ | $0.8689 \pm 0.00055$ | $0.8610 \pm 0.00056$ |
| Shape-Only | $0.5381 \pm 0.00112$ | $0.5515 \pm 0.00118$ | $0.5591 \pm 0.00122$ | $0.5530 \pm 0.00120$ |

The shape-only model shows a substantial drop in performance (macro-F1 = 0.5530), confirming that morphological descriptors alone are not sufficient for robust protocol in-ference. While morphology encodes meaningful geometric cues, these cues lack the fine-grained appearance information captured directly from images, explaining the large performance gap. The no-fusion model (macro-F1 = 0.8610) underperforms relative to the full model (0.8681), indicating that simply parallelizing the two modalities is less effective than an integrated fusion mechanism. This supports our design choice to enable deeper cross-modal interaction rather than treating the modalities independently.

Removing the transformer fusion block results in a small but consistent reduction in macro-F1 (0.8650), suggesting that the transformer refines cross-modal relationships, particularly when morphological features only partially align with visual appearance patterns. Overall, these results indicate that (1) image features carry the majority of predictive signal, (2)morphology provides complementary structural information, and (3) explicit fusion, especially transformer-based fusion, yields the most stable performance improvements.

## Conclusion

We introduced inverse protocol prediction (IPP) from single spheroid images, showing that bright-field morphology encodes recoverable signals of experimental conditions. Using SLiMIA, we evaluated segmentation, IPP, and temporal forecasting across convolutional, transformer, hybrid, feature-augmented, and hierarchical architectures. Hybrid models such as CoAtNet achieved the best accuracy–efficiency tradeoff, while fusion and hierarchical designs improved interpretability, structured consistency, and cross-domain robustness. Cross-dataset validation on RxRx1 and the Cell Tracking Challenge demonstrated that models incorporating explicit priors and structured dependencies generalize more effectively under domain shift, and that memory-enhanced recurrent models better retain temporal dynamics. Grad-CAM analysis confirmed reliance on biologically meaningful cues, while revealing sensitivity to dataset-specific artifacts. Per-label evaluation showed that strongly expressed morphological attributes are reliably recoverable, whereas weak or artifact-driven variables remain challenging. Future work should expand longitudinal depth, reduce technical biases, and integrate mechanistic priors with data-driven modeling to improve temporal fidelity and cross-dataset robustness. Together, our findings position protocol-aware, structured learning as a promising direction for interpretable and generalizable AI in experimental biology.

## Supplementary Note 1: Appendix

### A. Additional Results

#### A.1 Per-label Inverse Protocol Prediction (IPP) Performance Evaluation

We provide detailed per-label analyses to complement the aggregate results presented in the main text. Rather than presenting only overall accuracy, we focus on the label-wise F1 scores, which are more sensitive to class imbalance. Below, we summarize key observations.

**Cell Line, Culture Medium, and Formation Method:** These biologically central attributes were predicted with very high fidelity. The Image–Shape Fusion Transformer achieved the best F1 on cell line (0.9944), leveraging explicit morphological descriptors (e.g., circularity, eccentricity, axis lengths) that encode lineage-specific growth signatures beyond raw image texture. ViT-B/16 performed best on culture medium (F1 = 0.9642), benefiting from its global receptive field to capture medium-induced differences in spheroid compactness and necrotic core structure. However, these results partly reflect dataset imbalance: DMEMLG alone accounts for 38.9% of all images, with the top-3 media dominating overall. Finally, ConvNeXt achieved near-ceiling performance on formation method (F1 = 0.9949), as differences between agarose overlay, ULA plates, and hanging drops are morphologically striking, particularly at the spheroid boundary. Together, these labels highlight how explicit priors, transformer global context, and convolutional locality biases each confer complementary strengths depending on the task. **Seeding Density and Timepoint:** Both attributes proved more difficult due to long-tailed distributions and overlapping morphological ranges. CoAtNet achieved the highest F1 on seeding density (0.9302), with its hybrid convolution–attention design balancing local texture cues against global structure. Yet, densities such as 1000 vs. 2000 cells produce spheroids with similar compactness, limiting separability even for the best models. Timepoint prediction showed an even starker imbalance: although most models reached high accuracy (*>* 0.96), ConvNeXt-Tiny obtained the best F1 (0.8331). This discrepancy arises from extreme fragmentation—over 100 distinct timepoint values exist, but late stages dominate (e.g., 168 h = 25.4%, 96 h = 23.5%), while early and intermediate hours are sparse. As a result, models tend to over-predict majority late hours. Local convolutional features appear more robust to gradual morphological progression, explaining ConvNeXt’s advantage, though reframing timepoint as an ordinal or binned task may improve future performance.

**Biological Replicate:** Replicate-level prediction illustrates the limits of morphological inference. ConvNeXt achieved the best F1 on biological replicate (0.9583), likely due to its sensitivity to subtle texture differences between independent cultures.

**Microscope and Magnification:** Both attributes reached near-perfect performance across all models (*F* 1 *>* 0.999). These results, however, reflect dataset artifacts rather than biological signal: each microscope and magnification has distinct optical signatures such as field of view, resolution, and scale bar rendering. Models effectively memorize these acquisition-specific cues. While this provides a sanity check on model capacity, it should not be interpreted as evidence of meaningful biological inference.

In summary, attributes with strong morphological cues (cell line, culture medium, formation method) are predicted with high fidelity, while labels with fragmented or weak visual encoding (timepoint, seeding density) remain challenging. Biological replicate is moderately well captured, whereas microscope and magnification are trivially solved due to acquisition artifacts. These results highlight both the promise and the limitations of IPP: success depends as much on dataset balance and label definition as on model choice.

### B. Hyperparameters of Models

#### B.1 U-Net++

[leftmargin=1.5cm,style=nextline]

**Architecture** U-Net++ (UnetPlusPlus), encoder: ResNet-

50 (ImageNet), input channels: 3 (RGB), output classes: 1 (binary mask), activation: none (handled in loss/metrics)

**Loss** Focal Tversky Loss, *α* = 0.7, *β* = 0.3, *γ* = 0.75

**Optimizer** Adam, learning rate 1.0 *×* 10^*−*4^

**Scheduler** ReduceLROnPlateau (mode=max, patience=5, factor=0.5)

**Training** Up to 200 epochs, early stopping patience 40, batch size 8, shuffle train/validation, num workers 2, pin memory true

#### B.2 DeepLabV3

[leftmargin=1.5cm,style=nextline]

**Architecture** DeepLabV3 with ResNet-34 backbone (ImageNet), input 3 (RGB), output 1 (binary mask), activation: none

**Loss** 0.5*×* BCEWithLogits + 0.5*×* DiceLoss (Dice smooth 1 *×* 10^*−*6^)

**Optimizer** AdamW, learning rate 3.0 *×* 10^*−*4^, weight decay 1.0 *×* 10^*−*4^

**Scheduler** ReduceLROnPlateau (mode=max on validation Dice, patience=5, factor=0.5)

**Training** 200 epochs, early stopping 40, batch size 8, shuffle train, num workers 2, pin memory true

#### B.3 Attention U-Net

[leftmargin=1.5cm,style=nextline]

**Architecture** Attention U-Net, input 3 (RGB), output 1 (binary mask), activation: none

**Loss** BCEWithLogitsLoss + DiceLoss (0.5 each, Dice smooth 1.0 *×* 10^*−*6^)

**Optimizer** AdamW, learning rate 3.0 *×* 10^*−*4^

**Scheduler** ReduceLROnPlateau (mode=max on Dice Score, patience=5, factor=0.5)

**Training** 200 epochs, early stopping 40, batch size 8, shuffle train/validation, num workers 2, pin memory true

#### B.4 Swin-UNet

[leftmargin=1.5cm,style=nextline]

**Architecture** Swin-UNet (custom decoder), input 3 (RGB), output 1 (binary mask), backbone output features 768, decoder Conv–ReLU–Upsample stacks to 224*×*224

**Loss** BCEWithLogitsLoss

**Optimizer** Adam, learning rate 1.0 *×* 10^*−*4^

**Scheduler** ReduceLROnPlateau (mode=min, patience=5, factor=0.5)

**Mixed Precision** Enabled (torch.cuda.amp)

**Training** 200 epochs, early stopping 40, batch size 8, shuffle train, num workers 2, pin memory true

#### B.5 TransUNet

[leftmargin=1.5cm,style=nextline]

**Architecture** Transformer encoder + CNN decoder, input 3 (RGB), output 1 (binary mask), ViT output [B,196,768] reshaped to [B,768,14,14], decoder Conv2d–ReLU–Upsample (5 layers)

**Loss** BCEWithLogitsLoss

**Optimizer** Adam, learning rate 1.0 *×* 10^*−*4^

**Scheduler** ReduceLROnPlateau (mode=min, patience=5, factor=0.5)

**Mixed Precision** Enabled (torch.cuda.amp.autocast)

**Training** 200 epochs, early stopping 40, batch size 8, shuffle train, num workers 2, pin memory true

#### B.6 DeepLabV3+

[leftmargin=1.5cm,style=nextline]

**Architecture** DeepLabV3+ with ResNet-50 backbone (ImageNet), input 3 (RGB), output 1 (binary mask), activation: none

**Loss** Focal Tversky Loss, *α* = 0.7, *β* = 0.3, *γ* = 0.75

**Optimizer** Adam, learning rate 1.0 *×* 10^*−*4^

**Scheduler** ReduceLROnPlateau (mode=max on Dice, patience=5, factor=0.5)

**Mixed Precision** Enabled (torch.cuda.amp)

**Training** 200 epochs, early stopping 40, batch size 8, shuffle train/validation, num workers 2, pin memory true

#### B.7 RefineNet

[leftmargin=1.5cm,style=nextline]

**Architecture** RefineNet with ResNet-34 backbone (ImageNet), input 3 (RGB), output 1 (binary mask), activation: none

**Loss** Focal Tversky Loss, *α* = 0.7, *β* = 0.3, *γ* = 0.75

**Optimizer** Adam, learning rate 1.0 *×* 10^*−*4^

**Scheduler** ReduceLROnPlateau (mode=max on Dice Score, patience=5, factor=0.5)

**Mixed Precision** Disabled

**Training** 200 epochs, early stopping 40, batch size 8, shuffle train/validation, num workers 2, pin memory true

**Data** Input size 256*×* 256, augmentations: HorizontalFlip 0.5, VerticalFlip 0.3, RandomBrightnessContrast 0.3,

RandomGamma 0.3, Resize 256*×*256

#### B.8 SegNet

[leftmargin=1.5cm,style=nextline]

**Architecture** SegNet (VGG-style conv blocks), input 3 (RGB), output 1 (binary mask), activation: none

**Loss** Focal Tversky Loss, *α* = 0.7, *β* = 0.3, *γ* = 0.75

**Optimizer** Adam, learning rate 1.0 *×* 10^*−*4^

**Scheduler** ReduceLROnPlateau (mode=max on Dice Score, patience=5, factor=0.5)

**Mixed Precision** Enabled (torch.cuda.amp)

**Training** 200 epochs, early stopping 40, batch size 8, shuffle train/validation, num workers 2, pin memory true

**Data** Input size 256 *×*256, augmentations: HorizontalFlip 0.5, VerticalFlip 0.3, RandomBrightnessContrast 0.3, RandomGamma 0.3, GaussNoise, ElasticTransform, GridDistortion, ShiftScaleRotate, Resize 256*×*256

#### B.9 ConvNeXt-Tiny

[leftmargin=1.6cm,style=nextline]

**Model** ConvNeXt-Tiny backbone + multi-head classifier (one head per label)

**Pretrained Weights** ImageNet

**Output Heads** 9 (microscope, cell_line, culture_medium, formation_method, seeding_density, timepoint, biological_rep, technical_rep, magnification)

**Input** 224*×*224 RGB images

**Loss** CrossEntropyLoss (separate for each head)

**Optimizer** Adam, learning rate 1.0*×*10^*−*4^

**Scheduler** ReduceLROnPlateau (mode=max, patience=5)

**Training** up to 200 epochs, early stopping patience 40, batch size 32, shuffle (train only), 2 workers, pin memory

**Data** Train 6,418 images (T1–T8), val 714 images (T1–T8), test 7,999 images (T1–T24 unseen)

**Transforms (train)** Resize 224 *×*224, RandomHorizontalFlip, RandomRotation(10^*°*^), normalization mean=std=[0.5]

**Transforms (val)** Resize 224 *×*224, normalization mean=std=[0.5]

#### B.10 ViT-B/16

[leftmargin=1.6cm,style=nextline]

**Model** Vision Transformer (vit_base_patch16_224, timm pretrained)

**Input** 224*×*224 RGB images

**Output Heads** 9 (one per label column)

**Loss** Multi-task CrossEntropyLoss (sum over tasks)

**Optimizer** Adam, learning rate 1.0*×*10^*−*4^

**Scheduler** ReduceLROnPlateau (mode=max, patience=5, factor=0.5)

**Training** 200 epochs, early stopping 40, batch size 32, shuffle (train only), 2 workers, pin memory

**Data** Train 6,418 images (T1–T8), val 714 images (T1–T8), test 7,999 images (T1–T24 unseen)

**Transforms** Same as ConvNeXt

#### B.11 CoAtNet-0

[leftmargin=1.6cm,style=nextline]

**Model** CoAtNet-0 (timm coatnet_0_224, not pretrained)

**Input** 224*×*224 RGB images **Output Heads** 9 (one per label) **Loss** Multi-task CrossEntropyLoss

**Optimizer** Adam, learning rate 1.0*×*10^*−*4^

**Scheduler** ReduceLROnPlateau (mode=max, patience=5, factor=0.5)

**Training** 200 epochs, early stopping 40, batch size 32, 2 workers, pin memory

**Data** Train 6,418 images (T1–T8), val 714 images (T1–T8), test 7,999 images (T1–T24 unseen)

**Transforms** Same as ConvNeXt

#### B.12 Image Shape Fusion Transformer

[leftmargin=1.6cm,style=nextline]

**Model** ConvNeXt-Tiny backbone (ImageNet pretrained) + shape features projected as tokens fused via Transformer Encoder + multi-head classifier

**Fusion** Image token + 9 shape tokens *→*Transformer Encoder

**Transformer** *d*_*model*_=256, n_heads=4, n_layers=3, feedforward dim=512, dropout=0.1

**Shape Features** 9 geometric features (area, perimeter, eccentricity, solidity, extent, equivalent_diameter, major_axis_length, minor_axis_length, circularity), z-normalized

**Loss** Class-weighted CrossEntropyLoss

**Optimizer** AdamW, learning rate 1.0*×*10^*−*4^, weight decay

1.0*×*10^*−*2^

**Scheduler** ReduceLROnPlateau (mode=max on validation macro-F1, patience=5)

**Training** up to 200 epochs, early stopping patience 40, batch size 32, 4 workers, pin memory

**Data** Train 6,418 images (T1–T8), val 714 images (T1–T8), test 7,999 images (T1–T24 unseen)

**Transforms** Same resizing/augmentation as ConvNeXt

#### B.13 Hierarchical Multi-Task Transformer (HMTT)

[leftmargin=1.6cm,style=nextline]

**Model** ViT-B/16 encoder (ImageNet pretrained) + hierarchical multi-task heads conditioned on causal label order

**Label Order** cell_line *→* culture_medium *→* seeding_density *→* magnification *→* microscope *→* timepoint *→* biological_rep *→* technical_rep

**Output Heads** 8 (one per label)

**Loss** CrossEntropyLoss with class weights; optional Focal Loss (*γ*=2.0, *α*=0.25)

**Optimizer** AdamW, learning rate 1.0*×*10^*−*4^, weight decay 1.0*×*10^*−*2^

**Scheduler** ReduceLROnPlateau (mode=max validation macro-F1, patience=5)

**Training** up to 200 epochs, early stopping 40, batch size 32, 2 workers, pin memory, mixed precision (AMP)

**Data** Train 6,418 images (T1–T8), val 714 images (T1–T8), test 7,999 images (T1–T24 unseen)

**Transforms** Same as ConvNeXt

#### B. 14 ConvLSTM

[leftmargin=1.6cm,style=nextline]

**Data** Image size 128; sequence length 2; min gap 1; max gap 3; train/val/test split 70/15/15

**Training** Batch size 8; 4 workers; 200 epochs; learning rate 1.0 *×*10^*−*4^; optimizer Adam; loss L1; shuffle (train) True, (val/test) False; no gradient clipping or scheduler; mixed precision disabled; early stopping metric SSIM with patience 40

**Model** ConvLSTM architecture; hidden dimension 32; kernel size 3; decoder Conv2d *→* 1 channel

#### B.15 PredRNN++

[leftmargin=1.6cm,style=nextline]

**Data** Image size 128; sequence length 2; min frame gap 1; max frame gap 3; dataset split 70/15/15; batch size 8; 4 workers

**Model** PredRNN++ with two SpatioTemporal LSTM layers

+ Conv decoder; input channels 1; hidden dimensions [32, 64]; kernel size 3; decoder Conv2d (64*→* 1, kernel 1)

**Training** 200 epochs; optimizer Adam; learning rate 1.0 *×*10^*−*4^; loss L1; early stopping patience 40 (on SSIM);

#### B.16 MMFusionNet

[leftmargin=1.6cm,style=nextline]

**Data** Image size 128; sequence length 2; min gap 1; max gap 3; train/val/test split 70/15/15

**Model** Base channels 32; encoder 4 ConvBlocks with pooling; bottleneck 256 channels; decoder symmetric upsampling with skip connections; output 1-channel (sigmoid); categorical embedding dim 32; continuous features 2 (seeding density, timepoint); metadata fusion MLP (256*→*256) + FiLM conditioning

**Training** Batch size 8; 4 workers; 200 epochs; learning rate 1.0 *×*10^*−*4^; optimizer AdamW; scheduler ReduceLROnPlateau (patience 10, factor 0.5); loss *×*0.8 MSE + 0.2*×* (1–SSIM) if MS-SSIM available, else 0.6 *×*MSE + 0.4 *×*L1; early stopping patience 40; visual samples saved every 5 epochs; shuffle train=True, val=False

#### B.17 PhyDNet

[leftmargin=1.6cm,style=nextline]

**Data** Image size 128; context frames 2; prediction steps 1; frame gap 1–3; dataset split 70/15/15; batch size 8; 4 workers; seed 42

**Model** PhyDNet encoder + PhyCell + ConvLSTM residual + decoder; input channels 1; encoder channels 64; physics channels 32; residual channels 32; physics operators per channel 3; physics kernel size 3

**Training** 200 epochs; optimizer Adam; learning rate 1.0 *×*10^*−*4^; loss L1; mixed precision enabled (AMP); scheduler ReduceLROnPlateau (factor 0.5, patience 8); early stopping patience 40 on SSIM; best model

## Bibliography

1. Aliya Fatehullah, Si Hui Tan, and Nick Barker. Organoids as an in vitro model of human development and disease. Nature cell biology, 18(3):246–254, 2016.

2. Maria Chatzinikolaidou. Cell spheroids: the new frontiers in in vitro models for cancer drug validation. Drug discovery today, 21(9):1553–1560, 2016.

3. Rasheena Edmondson, Jessica Jenkins Broglie, Audrey F Adcock, and Liju Yang. Threedimensional cell culture systems and their applications in drug discovery and cell-based biosensors. Assay and drug development technologies, 12(4):207–218, 2014.

4. Taeyun Park, Taeyul K Kim, Yoon Dae Han, Kyung-A Kim, Hwiyoung Kim, and Han Sang Kim. Development of a deep learning based image processing tool for enhanced organoid analysis. Scientific reports, 13(1):19841, 2023.

5. Zhichao Liu, Luhong Jin, Jincheng Chen, Qiuyu Fang, Sergey Ablameyko, Zhaozheng Yin, and Yingke Xu. A survey on applications of deep learning in microscopy image analysis. Computers in biology and medicine, 134:104523, 2021.

6. Rabeea Fatma Khan, Byoung-Dai Lee, and Mu Sook Lee. Transformers in medical image segmentation: a narrative review. Quantitative Imaging in Medicine and Surgery, 13(12): 8747, 2023.

7. Simon Reiß, Constantin Seibold, Alexander Freytag, Erik Rodner, and Rainer Stiefelhagen. Every annotation counts: Multi-label deep supervision for medical image segmentation. In Proceedings of the IEEE/CVF conference on computer vision and pattern recognition, pages 9532–9542, 2021.

8. Kai Han, Victor S Sheng, Yuqing Song, Yi Liu, Chengjian Qiu, Siqi Ma, and Zhe Liu. Deep semi-supervised learning for medical image segmentation: A review. Expert Systems with Applications, 245:123052, 2024.

9. Lennart R Koetzier, Jie Wu, Domenico Mastrodicasa, Aline Lutz, Matthew Chung, W Adam Koszek, Jayanth Pratap, Akshay S Chaudhari, Pranav Rajpurkar, Matthew P Lungren, et al. Generating synthetic data for medical imaging. Radiology, 312(3):e232471, 2024.

10. Jiequan Zhang, Qingyu Zhao, Ehsan Adeli, Adolf Pfefferbaum, Edith V Sullivan, Robert Paul, Victor Valcour, and Kilian M Pohl. Multi-label, multi-domain learning identifies compounding effects of hiv and cognitive impairment. Medical image analysis, 75:102246, 2022.

11. Sumit Madan, Manuel Lentzen, Johannes Brandt, Daniel Rueckert, Martin Hofmann-Apitius, and Holger Fröhlich. Transformer models in biomedicine. BMC medical informatics and decision making, 24(1):214, 2024.

12. Raj Dave, Kshipra Pandey, Ritu Patel, Nidhi Gour, and Dhiraj Bhatia. Leveraging 3d cell culture and ai technologies for next-generation drug discovery. Cell Biomaterials, 1(3), 2025.

13. Alfredo De Cillis, Valeria Garzarelli, Alessia Foscarini, Giuseppe Gigli, Antonio Turco, Elisabetta Primiceri, Maria Serena Chiriacò, and Francesco Ferrara. 3d-printed barriers with machine learning powered image analysis for enhanced wound healing assays. Materials & Design, page 114746, 2025.

14. Songshan Zhu, Jun Yin, Xiaotong Lu, Dan Jiang, Rui Chen, Kai Cui, Wanjun He, Na Huang, and Guangxian Xu. Influence of experimental variables on spheroid attributes. Scientific Reports, 15(1):9751, 2025.

15. Isabel Kemmer, Antje Keppler, Beatriz Serrano-Solano, Arina Rybina, Buğra Özdemir, Johanna Bischof, Ayoub El Ghadraoui, John E Eriksson, and Aastha Mathur. Building a fair image data ecosystem for microscopy communities. Histochemistry and Cell Biology, 160 (3):199–209, 2023.

16. Maximiliaan Huisman, Mathias Hammer, Alex Rigano, Ulrike Boehm, James J Chambers, Nathalie Gaudreault, Alison J North, Jaime A Pimentel, Damir Sudar, Peter Bajcsy, et al. A perspective on microscopy metadata: data provenance and quality control. arXiv preprint arXiv:1910.11370, 2019.

17. Paula Montero Llopis, Rebecca A Senft, Tim J Ross-Elliott, Ryan Stephansky, Daniel P Keeley, Preman Koshar, Guillermo Marqués, Ya-Sheng Gao, Benjamin R Carlson, Thomas Pengo, et al. Best practices and tools for reporting reproducible fluorescence microscopy methods. Nature Methods, 18(12):1463–1476, 2021.

18. Ugis Sarkans, Wah Chiu, Lucy Collinson, Michele C Darrow, Jan Ellenberg, David Grunwald, Jean-Karim Hériché, Andrii Iudin, Gabriel G Martins, Terry Meehan, et al. Rembi: Recommended metadata for biological images—enabling reuse of microscopy data in biology. Nature methods, 18(12):1418–1422, 2021.

19. Eva Blondeel, Arne Peirsman, Stephanie Vermeulen, Filippo Piccinini, Felix De Vuyst, Diogo Estêvão, Sayida Al-Jamei, Martina Bedeschi, Gastone Castellani, Tânia Cruz, et al. The spheroid light microscopy image atlas for morphometrical analysis of three-dimensional cell cultures. Scientific Data, 12(1):283, 2025.

20. Kingma DP Ba J Adam et al. A method for stochastic optimization. arXiv preprint arXiv:1412.6980, 1412(6), 2014.

21. Ilya Loshchilov and Frank Hutter. Decoupled weight decay regularization. arXiv preprint arXiv:1711.05101, 2017.

22. Leslie N Smith. Cyclical learning rates for training neural networks. In 2017 IEEE winter conference on applications of computer vision (WACV), pages 464–472. IEEE, 2017.

23. Seyed Sadegh Mohseni Salehi, Deniz Erdogmus, and Ali Gholipour. Tversky loss function for image segmentation using 3d fully convolutional deep networks. In International workshop on machine learning in medical imaging, pages 379–387. Springer, 2017.

24. Fausto Milletari, Nassir Navab, and Seyed-Ahmad Ahmadi. V-net: Fully convolutional neural networks for volumetric medical image segmentation. In 2016 fourth international conference on 3D vision (3DV), pages 565–571. Ieee, 2016.

25. Zongwei Zhou, Md Mahfuzur Rahman Siddiquee, Nima Tajbakhsh, and Jianming Liang. Unet++: A nested u-net architecture for medical image segmentation. In International workshop on deep learning in medical image analysis, pages 3–11. Springer, 2018.

26. Liang-Chieh Chen, Yukun Zhu, George Papandreou, Florian Schroff, and Hartwig Adam. Encoder-decoder with atrous separable convolution for semantic image segmentation. In Proceedings of the European conference on computer vision (ECCV), pages 801–818, 2018.

27. Ozan Oktay, Jo Schlemper, Loic Le Folgoc, Matthew Lee, Mattias Heinrich, Kazunari Misawa, Kensaku Mori, Steven McDonagh, Nils Y Hammerla, Bernhard Kainz, et al. Attention u-net: Learning where to look for the pancreas. arXiv preprint arXiv:1804.03999, 2018.

28. Hu Cao, Yueyue Wang, Joy Chen, Dongsheng Jiang, Xiaopeng Zhang, Qi Tian, and Manning Wang. Swin-unet: Unet-like pure transformer for medical image segmentation. In European conference on computer vision, pages 205–218. Springer, 2022.

29. Ian Goodfellow. Deep learning, 2016.

30. Jieneng Chen, Yongyi Lu, Qihang Yu, Xiangde Luo, Ehsan Adeli, Yan Wang, L. Lu, Alan L Yuille, and Yuyin Zhou. Transunet: Transformers make strong encoders for medical image segmentation. arXiv preprint arXiv:2102.04306, 2021.

31. Guosheng Lin, Anton Milan, Chunhua Shen, and Ian Reid. Refinenet: Multi-path refinement networks for high-resolution semantic segmentation. In Proceedings of the IEEE conference on computer vision and pattern recognition, pages 1925–1934, 2017.

32. Vijay Badrinarayanan, Alex Kendall, and Roberto Cipolla. Segnet: A deep convolutional encoder-decoder architecture for image segmentation. IEEE transactions on pattern analysis and machine intelligence, 39(12):2481–2495, 2017.

33. Zhuang Liu, Hanzi Mao, Chao-Yuan Wu, Christoph Feichtenhofer, Trevor Darrell, and Saining Xie. A convnet for the 2020s. In Proceedings of the IEEE/CVF conference on computer vision and pattern recognition, pages 11976–11986, 2022.

34. Abdullah A Asiri, Ahmad Shaf, Tariq Ali, Muhammad Ahmad Pasha, Muhammad Aamir, Muhammad Irfan, Saeed Alqahtani, Ahmad Joman Alghamdi, Ali H Alghamdi, Abdullah Fahad A Alshamrani, et al. Advancing brain tumor classification through fine-tuned vision transformers: A comparative study of pre-trained models. Sensors, 23(18):7913, 2023.

35. Zihang Dai, Hanxiao Liu, Quoc V Le, and Mingxing Tan. Coatnet: Marrying convolution and attention for all data sizes. Advances in neural information processing systems, 34: 3965–3977, 2021.

36. Jingsong Xia, Yue Yin, and Xiuhan Li. An efficient medical image classification method based on a lightweight improved convnext-tiny architecture. arXiv preprint arXiv:2508.11532, 2025.

37. Bardia Rafieian and Pere-Pau Vázquez. Improved multi-label hierarchical patent classification using llms. World Patent Information, 81:102356, 2025.

38. Tsung-Yi Lin, Priya Goyal, Ross Girshick, Kaiming He, and Piotr Dollár. Focal loss for dense object detection. In Proceedings of the IEEE international conference on computer vision, pages 2980–2988, 2017.

39. Ling Zhang, L. Lu, Xiaosong Wang, Robert M Zhu, Mohammadhadi Bagheri, Ronald M Summers, and Jianhua Yao. Spatio-temporal convolutional lstms for tumor growth prediction by learning 4d longitudinal patient data. IEEE transactions on medical imaging, 39(4):1114–1126, 2019.

40. Yunbo Wang, Zhifeng Gao, Mingsheng Long, Jianmin Wang, and Philip S Yu. Predrnn++: Towards a resolution of the deep-in-time dilemma in spatiotemporal predictive learning. In International conference on machine learning, pages 5123–5132. PMLR, 2018.

41. Oliver Schön, Ricarda-Samantha Götte, and Julia Timmermann. Multi-objective physicsguided recurrent neural networks for identifying non-autonomous dynamical systems. IFAC-PapersOnLine, 55(12):19–24, 2022.

42. Dominik Klein, Giovanni Palla, Marius Lange, Michal Klein, Zoe Piran, Manuel Gander, Laetitia Meng-Papaxanthos, Michael Sterr, Lama Saber, Changying Jing, et al. Mapping cells through time and space with moscot. Nature, 638(8052):1065–1075, 2025.

43. Ethan Perez, Florian Strub, Harm De Vries, Vincent Dumoulin, and Aaron Courville. Film: Visual reasoning with a general conditioning layer. In Proceedings of the AAAI conference on artificial intelligence, volume 32, 2018.

44. Vincent Le Guen and Nicolas Thome. Disentangling physical dynamics from unknown factors for unsupervised video prediction. In Proceedings of the IEEE/CVF conference on computer vision and pattern recognition, pages 11474–11484, 2020.

45. Xiaowei Jia, Anuj Karpatne, Jared Willard, Michael Steinbach, Jordan Read, Paul C Hanson, Hilary A Dugan, and Vipin Kumar. Physics guided recurrent neural networks for modeling dynamical systems: Application to monitoring water temperature and quality in lakes. arXiv preprint arXiv:1810.02880, 2018.

46. Liang-Chieh Chen, Yukun Zhu, George Papandreou, Florian Schroff, and Hartwig Adam. Encoder-decoder with atrous separable convolution for semantic image segmentation. In Proceedings of the European conference on computer vision (ECCV), pages 801–818, 2018.

47. Fei Luo, Daoqi Wu, Luis Rojas Pino, and Weichao Ding. A novel multimodel medical image fusion framework with edge enhancement and cross-scale transformer. Scientific Reports, 15(1):11657, 2025.

48. Ramprasaath R Selvaraju, Michael Cogswell, Abhishek Das, Ramakrishna Vedantam, Devi Parikh, and Dhruv Batra. Grad-cam: Visual explanations from deep networks via gradient-based localization. In Proceedings of the IEEE international conference on computer vision, pages 618–626, 2017.

49. Aditya Chattopadhay, Anirban Sarkar, Prantik Howlader, and Vineeth N Balasubramanian. Grad-cam++: Generalized gradient-based visual explanations for deep convolutional networks. In 2018 IEEE winter conference on applications of computer vision (WACV), pages 839–847. IEEE, 2018.

50. Maciej Sypetkowski, Morteza Rezanejad, Saber Saberian, Oren Kraus, John Urbanik, James Taylor, Ben Mabey, Mason Victors, Jason Yosinski, Alborz Rezazadeh Sereshkeh, et al. Rxrx1: A dataset for evaluating experimental batch correction methods. In Proceedings of the IEEE/CVF conference on computer vision and pattern recognition, pages 4285–4294, 2023.

51. Cell tracking challenge. http://celltrackingchallenge.net/.

